# Identification of *TTN* as a novel candidate gene for atrioventricular block in a Chinese pedigree by whole-exome sequencing

**DOI:** 10.1101/506824

**Authors:** Guohui Liu, Ziying Yang, Weiwei Chen, Junguang Xu, Liangwei Mao, Qinlin Yu, Jian Guo, Hui Xu, Fengxia Liu, Yan Sun, Hui Huang, Zhiyu Peng, Jun Sun, Wei Li, Ping Yang

**Affiliations:** Department of Cardiology, China-Japan Union Hospital, Jilin University, Changchun 100029, Jilin Province, China; Tianjin Medical Laboratory, BGI-Tianjin, BGI-Shenzhen, Tianjin 300308, China; Binhai Genomics Institute, BGI-Tianjin, BGI-Shenzhen, Tianjin 300308, China; James D. Watson Institute of Genome Sciences, Hangzhou 310058, China; BGI Genomics, BGI-Shenzhen, Shenzhen 518083, China; Department of Molecular Cell Biology, UC Berkeley, Berkeley, CA 94704, USA

**Author notes:** Correspondence: Wei Li,; Ping Yang.

**Keywords:** autosomal dominant inheritance, atrioventricular block, *TTN*, whole-exome sequencing

## Abstract

**Purpose:** Cardiovascular diseases are the most common cause of death globally. In which atrioventricular block (AVB) is a common disorder with genetic causes, but the responsible genes have not been fully identified yet. To determine the underlying causative genes involved in cardiac AVB, here we report a three-generation Chinese family with severe autosomal dominant cardiac AVB that has been ruled out as being caused by known genes mutations.

**Methods:** Whole-exome sequencing was performed in five affected family members across three generations, and co-segregation analysis was validated on other members of this family.

**Results:** Whole-exome sequencing and subsequent co-segregation validation identified a novel germline heterozygous point missense mutation, c.49287C>A (p.N16429K), in the titin (TTN, NM_001267550.2) gene in all 5 affected family members, but not in the unaffected family members. The point mutation is predicted to be functionally deleterious by *in-silico* software tools. Our finding was further supported by the conservative analysis across species.

**Conclusion:** Based on this study, *TTN* was identified as a potential novel candidate gene for autosomal dominant AVB; this study expands the mutational spectrum of *TTN* gene and is the first to implicate *TTN* mutations as AVB disease causing in a Chinese pedigree.

## Introduction

Atrioventricular block (AVB) is a systemic disease characterized by the heart conduction block between the atria and the ventricles. The degrees of AVB are classified into three tiers according to the severity of the conduction impairment between the sinoatrial node and atrioventricular nodes.^1,2^ Clinical diagnosis for AVB is mainly determined by the electrocardiographs (ECG). The first degree of AVB, featured with a PR interval higher than 0.2 second, is typically asymptomatic, whereas the third degree of AVB is the most severe type and has serious symptoms including, but not limited, dizziness, fatigue, and syncope.^3^ Third degree of AVB has no association between P waves and QRS complexes, and usually the auricular rate is higher than the ventricular rate. A pacemaker implement is usually needed for the third degree AVB patient.^4,5^ AVB can result from various causes including ischemia, cardiomyopathy, fibrosis or drugs.^6–8^ The prevalence of third degree AVB is about 0.04% as reported previously.^9^

The cardiac conduction system consists of gap junction proteins and ion channels on the cell membrane. As reported previously, many gap junction and ion channel genes such as *KCNQ1, KCNE1, KCNH2, KCNE2, KCNJ2, SCN5A, CASQ2, ANK2, GJA5* and *HCN4* were known to relate with AVB.^10–16^ Besides, the structural protein TTN mutation (TTN p.Glu5365Asp;TTN p.Arg3067His; TTN p.Arg8985Cys) was reported previously to be detected among AVB patients.^6^ *TTN* mutation was reported previously to be the genetic cause of cardiomyopathy,^17,18^ one of the potential causes of AVB. Recently, *TTN* was identified as being associated with atrial and atrioventricular electrical activity though genome-wide association study.^19^ However, the relationship between *TTN* and AVB is illusive; the underlying genetic mechanism contributing to the third degree AVB is still not fully characterized.^1,20^

For rare and undiagnosed Mendelian diseases, whole exome sequencing (WES) is a good way to analyze coding regions among thousands of genes simultaneously through next generation sequencing method.^21^ WES has an overall molecular diagnostic rate of 25% to 50%,^22,23^ much higher than that of some other genetic tests including chromosomal microarray analysis (15%-20%)^24^ and chromosome tests (5%-10%).^25^ WES can even hit a 68% diagnostic rate when the iterative approach was performed.^26^ In this study, WES was performed to identify the underlying genetic cause of AVB among the pedigree according to the procedure reported previously.^27,28^

## Materials and Methods

### Information of the pedigree

The pedigree of a three-generation Chinese family with AVB is presented in **Fig 1A** and seems an autosomal dominant inheritance. The venous blood samples of five affected and two unaffected family members were collected for this study. Electrocardiographs (ECG) of the proband (affected) and his son (unaffected) are presented in **Fig 1B** and **Fig 1C**, respectively. This study was approved by the Institutional Review Board on Bioethics and Biosafety of BGI and all the 7 participants signed the informed consent.

**Fig 1.**
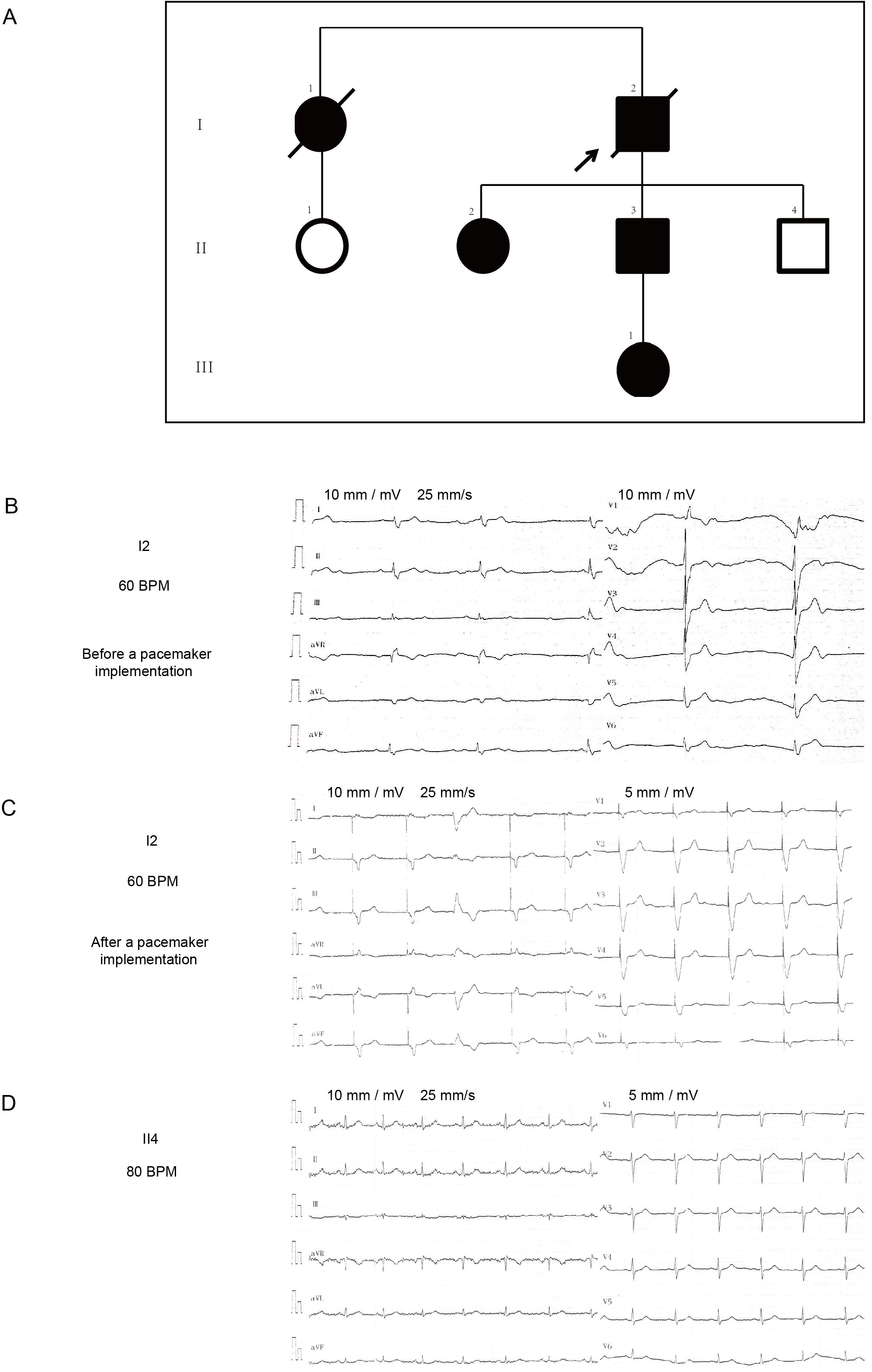
Pedigree and ECG photographs. (A) AVB pedigree of the three generation Chinese family. Full symbols represent affected members with AVB phenotypes, and open symbols represent members with normal phenotype. Squares and circles represent male and female members, respectively. The slash indicates the deceased member. The arrow represents the proband. All members shown have their venous blood samples collected. (B) ECG photographs of the proband before a pacemaker implementation. (C) ECG photographs of the proband after a pacemaker implementation. (D) ECG photographs of the proband’s son. BPM, beats per minute.

### Exome sequencing protocol

Five milliliter of peripheral blood was taken from each affected family member into an EDTA tube. Genomic DNA was extracted with a QIAamp DNA Blood Mini Kit (Qiagen, Germany). The DNA integrity was examined via the gel electrophoresis. One microgram genomic DNA was fragmented randomly to an average size of 250 base pair (bp) through the Covaris Acoustic System. After end-repairing and adenylation, adapters were ligated to the ends of the DNA fragments. The products were then amplified by ligation-mediated polymerase chain reaction (LM-PCR), and then hybridized with Roche 64 M whole exome capture kit to capture the target regions. Quantitative PCR was performed to evaluate the enrichment. Then the qualified library was sequenced on Hiseq2500 (Illumina Inc., San Diego, CA, USA) for 100 bp paired-end run.

### Reads alignment, variant calling, annotation

Illumina Pipeline (version 1.3.4) was used for generating the primary sequence data. The clean data containing pair-end reads was mapped to human genome (NCBI37/hg19) using BWA software (Burrows Wheeler Aligner http://sourceforge.net/projects/bio-bwa/). SNVs (single nucleotide variants) were called by using SOAP SNP software (http://soap.genomics.org.cn/) and small InDels (insertions and deletions) were called by SAM tools Pileup software (http://sourceforge.net/projects/samtools/). The variants called out were annotated using Gaea, an annotation pipeline in-house developed by BGI, to provide information regarding their effect on protein, allele frequency in the general population database, the relation between the phenotype and genotype, and the *in-silico* tools prediction results. The public databases used to calculate the frequency in normal population include 1K genome database (http://www.1000genomes.org/), ESP6500 database, dbSNP database, and BGI local database, which includes whole exome data of 2087 normal subjects. The software tools used for non-synonymous functional predictions in the annotation pipeline includes MutationTaster,^29^ SIFT,^30^ PROVEAN,^31^ and Ens Condel.^32^ The data are available in the China Genebank Nucleotide Sequence Archive (https://db.cngb.org/cnsa, accession number CNP0000271).

### Sanger Sequencing

Sanger sequencing of all reported potential pathogenic variants was performed for the proband as well as the other family members as shown in **Figure 1A**. Primers were designed to amplify products ranging from 300 to 600 base pair in length harboring the site of interest. *TTN* primers are as followed: forward 5’-GCTGCCACCATCATCATCTG-3’; reverse 5’-ATGTTGTGGAGAGACGAGCA-3’. PCR amplifications were conducted in a 25 μl reaction volume containing 1x PCR buffer with 1.5 mM MgCl_2_, 200 μM each dNTP, 0.25 U Taq DNA polymerase, 1.5 μM primers and 50 ng genomic DNA. The PCR condition was 94 °C for 4 min, 35 cycles of 94 °C for 30 s, 59 °C for 30 s, and 72 °C for 45 s, and a final 72 °C extension for 10 min. The PCR products were purified and sequenced on an ABI3730xl sequencer. The Sanger chromatogram was analyzed by the Sequencing Analysis 5.2.

## Results

### Characteristic of the patients

The pedigree is shown in **Figure 1A**. This three generation Chinese family with five affected AVB patients and two unaffected members is from the north of China. The pattern of inheritance seems to be autosome dominant (AD).

The ECG photographs of the proband (I2) and his unaffected son (II4) are shown in **Figure 1B** and **Figure 1D** respectively. The proband is affected with typical three degree AVB as is betrayed in **Figure 1B** (before a pacemaker implementation) and in **Figure 1C** (after a pacemaker implementation), whereas one of his son is unaffected with AVB as shown in **Figure 1D**. As is shown in **Figure 1B**, a typical three degree AVB can be observed with the ECG features that the association between P waves and QRS complexes are unestablished, and the auricular rate is higher than the ventricular rate. **Figure 1C** and **Figure 1D** show a normal ECG feature.

The characteristics of all 7 family members are shown in **Table 1**, among them three patients (I1, I2, and II3) were implemented with a pacemaker to facilitate cardiac beating conduction from the inception date at not determined (N.D.), 47 and 38 years old respectively, two of them (I1 and I2) progressed into heart failure, while the other one (II3) has not yet shown signs of heart failure at 49 years old.

### Exome sequencing and potential pathogenic mutations

Generally, about 10 Gigabases raw data output were generated for each patient, providing an average of 101 million sequencing reads with a 100-bp length mapped to target region and a higher than 110-fold mean depth across the target region. Also, the average coverage at >10× read depth of the exome was 93%, which is within the expected coverage and depth for WES studies. An average of 24,441 variants were identified in the exomes of each patient.

After the frequency filtration, appropriate functional and inheritance pattern analysis for this pedigree, all known AVB causing genes were excluded as the pathogenic reason in this pedigree. Eventually a total of 11 variants from 10 genes were called as potential pathogenic variants associated with AVB in this pedigree with autosome recessive (AR) pattern or AD pattern (**Table 2**). To validate the variants identified by WES, Sanger sequencing was performed for both affected and unaffected family members (**Figure 1A**).

After the sanger sequencing validation among the whole pedigree, only *TTN*: c.49287C>A (p.N16429K) (**Figure 2A**) was determined to be segregated with AVB phenotype in the proband as well as in the other affected members in this pedigree, whereas absent in unaffected family members as well as healthy population controls, in an autosome dominant inheritance pattern. This amino acid mutation site is evolutionarily conservative across species spectrum as shown in Figure 2B, implying a potential important and indispensable role for the proper protein function and execution. Meanwhile, the *in-silico* software tools including MutationTaster, SIFT, and PROVEAN predict this mutation as deleterious for the correct protein function (**Table 2**).

**Fig 2.**
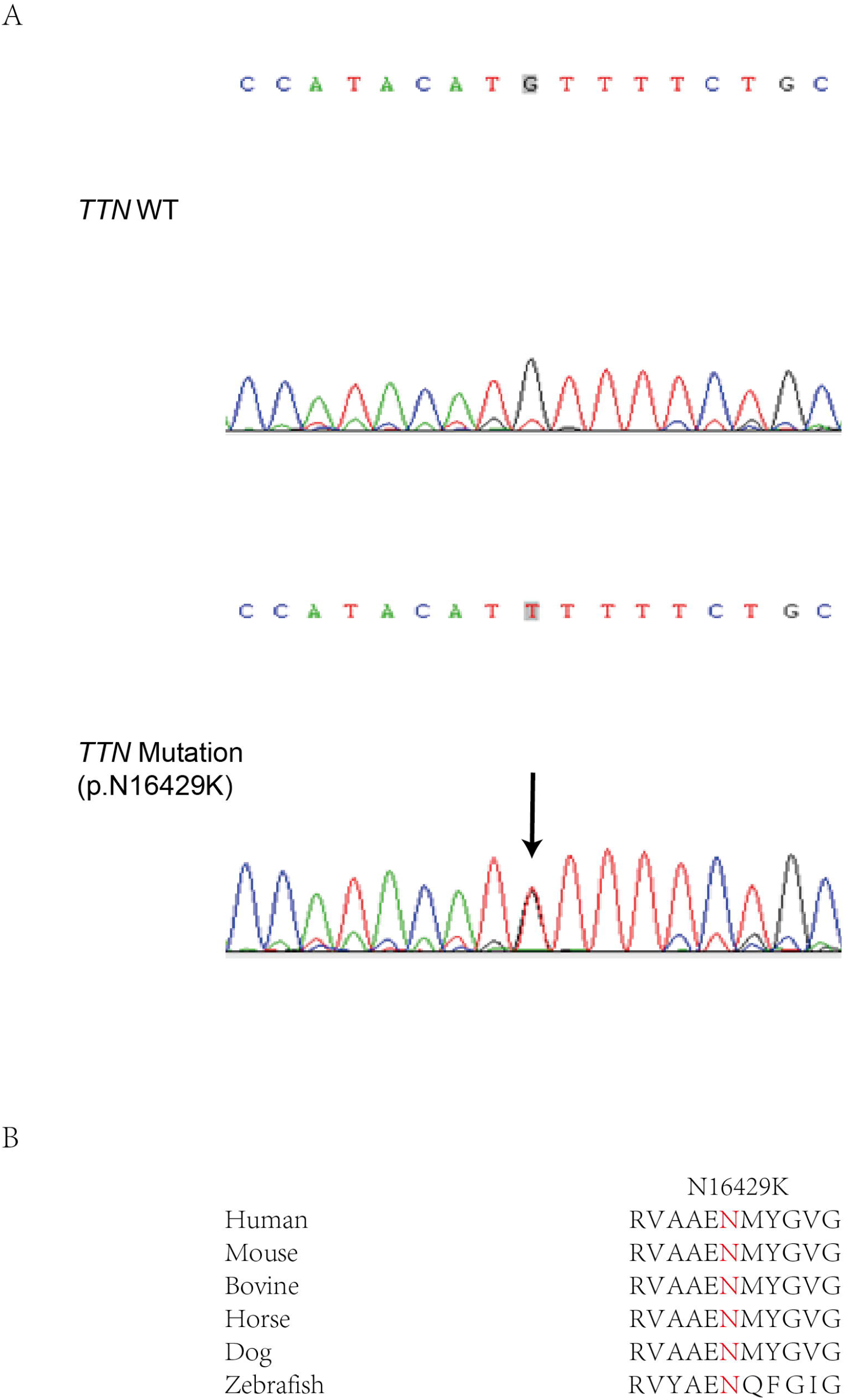
Identification and validation of the heterozygous pathogenic variant c.49287C>A in *TTN*. (A) Sequencing chromatogram of the wild type and the heterozygous c.49287C>A in *TTN*; (B) Schematic diagram of the conservativeness across species.

Taken together, according to the American College of Medical Genetics and Genomics (ACMG) guidelines issued in 2015^33^, this *TTN* mutation (*TTN*: c.49287C>A) can be interpreted as Likely Pathogenic (LP) which means greater than 90% certainty of this variant being disease-causing.

## Conclusion

Despite the fact that *TTN* has been reported to be associated with AVB, our study, for the first time, validates *TTN* as the disease-causing gene for AVB in a three-generation Chinese pedigree. The novel *TTN* gene variant c.49287C>A identified by WES is supportive for the genotype-phenotype correlation, and is co-segregated with AVB in this Chinese pedigree. Yet, the underlying mechanisms of *TTN* gene involved in AVB are for further exploration. Nevertheless, our new findings expand the mutational spectrum of *TTN* gene, provides evidence to understand the function of *TTN* gene in AVB, and shed light on the clinic diagnosis and potential treatment for the cardiac AVB disease.

## Acknowledgments

We would like to thank Jilin Province Development and Reform Commission (2016C026) for support.

We appreciate all patients and their families for their participation in this study.

## Author Contributions

All listed authors have each made substantial contributions to conception and design, acquisition of data, or analysis and interpretation of data; Wei Li participated in drafting the manuscript and revising it critically for content; Guohui Liu, Weiwei Chen and Ping Yang collected patient materials; Guohui Liu, Wei Li, Qinlin Yu and Ping Yang have approved the final version of the submitted manuscript. Guohui Liu, Ziying Yang and Ping Yang accepted responsibility for the integrity of the data analysis. All authors reviewed the manuscript.

---

**Table 1. The clinical characteristics and genetic testing results of the AV block pedigree.**

**Table 2. Summary of variants identified in the WES gene sequencing data.**

